# Characterization of cytotrophoblast cell population dynamics throughout pregnancy

**DOI:** 10.1101/2025.01.22.634235

**Authors:** Olivier Schäffers, Simone Verburgt, Madhavi Harhangi, Michelle Broekhuizen, Eline van Hengel, Lotte van der Meeren, Sam Schoenmakers, Dick Dekkers, Wouter Doff, Jeroen Demmers, Irwin Reiss, Joost Gribnau, Bas van Rijn

## Abstract

To understand cytotrophoblast (CTB) dynamics during pregnancy, we constructed a single-cell RNA sequencing dataset profiling over 35,000 CTBs isolated from human placental tissues at various gestational stages. Our analysis revealed a robust set of transcripts defining distinct CTB subsets and their spatial relationships within the placenta. We identified a transit-amplifying primitive trophoblast population that may contribute to the CTB pool and found that CTBs located at the smooth chorion form a leak-tight epithelial layer that adapts to low oxygen environments without undergoing syncytialization. Additionally, our study suggests that CTB fate in the villous chorion may be influenced by interactions with the basal lamina, as indicated by a population of BCAM+ CTBs. This subset, isolated from term placental tissue, displayed progenitor capabilities in organoid cultures, supporting the hypothesis that BCAM+ CTBs maintain a progenitor role late in pregnancy.

## Introduction

The placenta is a vital organ that develops during pregnancy, serving as the lifeline between the mother and the developing fetus. It is responsible for the exchange of nutrients, gases, and waste products, ensuring that the fetus receives the necessary sustenance for healthy growth (1). Central to the placenta’s function are the placental villi, which are tiny, finger-like projections extending from the chorionic surface into the maternal uterine tissue (2). Placental villi are classified into various types, such as stem villi, which provide structural support, and terminal villi, which are primarily involved in the exchange process. Post-conception, stem villi start to form around the embryo, which then progress asymmetrically to establish two distinct regions (3). At the implantation side, known as the embryonic pole, villi proliferate and branch out as terminal villi, forming the chorion frondosum (alternatively referred to as villous chorion). Conversely, stem villi degenerate at the abembryonic pole, resulting in the chorion laeve (or smooth chorion), which merges with the amnion to form the fetal membranes.

Structurally, placental villi are composed of multiple cell layers, including an outer layer of syncytiotrophoblasts (STBs) and an inner core of cytotrophoblasts (CTBs), and house a network of fetal blood vessels to create a large surface area for efficient nutrient and gas exchange. Placental villi continuously adapt and branch out throughout pregnancy, underscoring their essential role in maintaining fetal well-being and supporting the intricate balance of maternal-fetal interactions (4). This branching is facilitated by the interplay of various growth factors, signaling molecules, and extracellular matrix components that regulate trophoblast proliferation, differentiation, and migration (5). Additionally, the placental villi exhibit remarkable adaptability, responding to changes in maternal and fetal environments by modifying their structure and function (6, 7). Regeneration of damaged or aged villi ensures the continuous efficiency of the placenta, with mechanisms such as trophoblast turnover and angiogenesis playing crucial roles (8).

Currently, our understanding of the presence of putative trophoblast stem cell (TSC) populations in the human placenta is limited, with proliferative CTBs identified as trophoblast progenitor cells (5). These CTBs exhibit bipotentiality, capable of either fusing to form multinucleated STBs or migrating out of villi as extravillous trophoblasts (EVTs) to invade the uterus. Despite their importance, the molecular mechanisms guiding CTB fate throughout gestation remain partially understood, mainly due to the challenges in accessing and characterizing these cells at various developmental stages. In particular, questions continue to arise regarding CTB self-renewal and whether these cells persist or deplete throughout gestation. Previous studies have indicated that CTB stemness tends to decrease as pregnancy progresses, with proliferation significantly declining by the end of the first trimester (9, 10). A recent study, however, successfully generated trophoblast organoid cultures from term placental tissue, suggesting that CTBs maintain self-renewal capabilities till the end of pregnancy (11).

To better understand CTB dynamics during pregnancy, we constructed a single-cell RNA-seq dataset to dissect the transcriptional landscape of individual CTBs throughout early, mid, and late gestation. By profiling over 35,000 CTBs isolated from human placental tissues at distinct time points, we identified a robust set of transcripts that define subsets of CTB cells, including distinct spatial relationships within the placenta. Using embryo datasets, we found a transit-amplifying primitive trophoblast population that might give rise to the CTB pool. We reveal that CTBs located at the smooth chorion form a leak-tight epithelial layer that adapts to a low oxygen environment and do not undergo syncytialization.

Furthermore, we suggest that CTB fate in the villous chorion might be regulated by contact with the basal lamina, indicated by a population of BCAM^+^ CTBs. Our study is the first to comprehensively profile CTBs throughout pregnancy, shedding light on the dynamics underlying the development and adaptability of placental villi.

## Results

### CTB populations during gestation

To dissect the transcriptional landscape of CTBs along gestation, we analyzed publicly available scRNA-seq datasets of human pre-implantation and pre-gastrulation embryos (E3-E12/13) (12, 13), first trimester placentas (6-14 weeks) (14, 15), second trimester placentas (17-24 weeks) (16) and term placentas (39-40 weeks) (Figure 1A) (17). After quality control and data pre-processing, 110,683 cells (1954 cells derived from the embryo datasets; 108,729 cells derived from the placental datasets) were maintained for further analysis. We enriched for CTB populations using canonical markers. For the embryonic datasets, we first segregated the trophectoderm (TE) lineage, after which we selected *KRT7*^+^ *HLA-G*^-/dim^ *VIM*^-^ cells (Supplementary Figure 1). For the placental datasets, we directly selected *KRT7*^+^ *HLA-G*^-/dim^ *VIM*^-^ cells. Overall, we obtained 38,309 cells that after integration and low-resolution clustering could be grouped into four main populations (Figure 1B). As one population of CTBs was only found in the embryo datasets, we termed these cells TE-like CTB (Figure 1C). The other three CTB populations, smooth, villous and cycling CTB, were present in all gestational timepoints (Figure 1C). The relative number of cycling CTBs, along with their cell cycle activity, decreased over gestational time, but cycling CTB were still present in term placentas (Figure 1C, Supplementary Figure 2A). We stained for proliferative cells using Ki67 (*MKI67*), which is expressed in the entire cell cycle except G0 phase, and indeed detected Ki-67-positive cells in term placentas (Supplementary Figure 2B). This indicates that CTB populations with proliferative potential are still present in term placental tissue.

**Figure 1.**
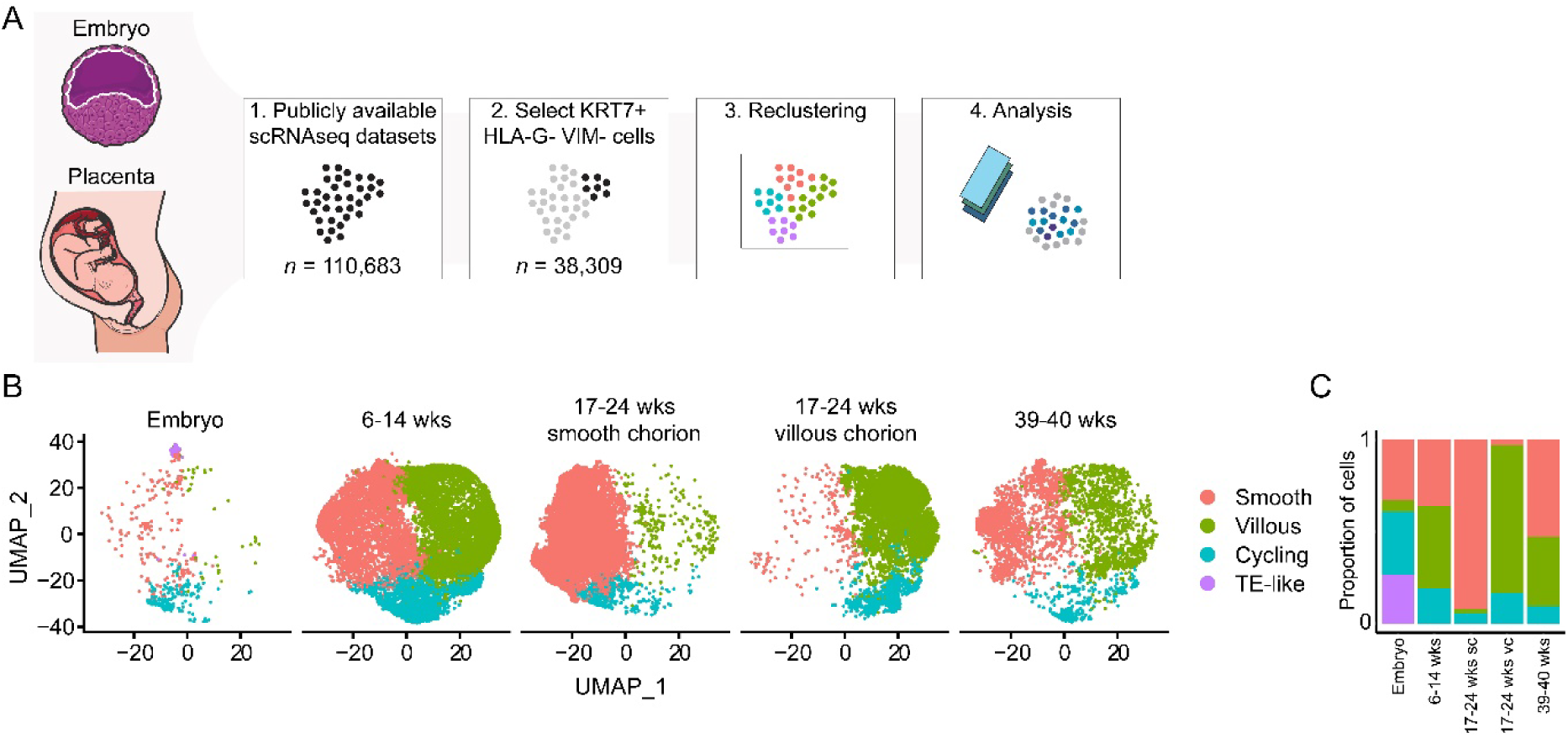
Single-cell transcriptional profiling of CTB subsets throughout gestation. (A) Schematic overview of our systematic approach to construct a CTB dataset using publicly available single-cell RNA-seq (scRNAseq) datasets. Datasets of human embryos and placental tissues were integrated and KRT7^+^ HLA-G^-^ VIM^-^ cells were selected to enrich for CTBs for downstream analyses. (B) UMAP plot showing 4 main clusters of CTB cells after integration of CTBs from embryo datasets (*n* = 674), first trimester placenta (*n* = 14,134), second trimester smooth and villous chorion (*n* = 20,048) and term placenta (*n* = 3453). (C) Proportion of CTB clusters for each gestational timepoint.

### TE-like CTB correlate to trophoblast stem cells

Since the TE-like CTB population appeared solely in the embryo datasets, we speculated whether it might represent a subset of primitive trophoblast cells. TE-like CTB showed enrichment for *EPCAM*, *GATA2* and *LAMA1* compared to all other CTBs, and depletion of *FN1* and *EGFR* (Figure 2A). Functional enrichment analysis showed that TE-like CTB differentially express transcripts related to adaptation to hypoxia, including anaerobic metabolism and HIF-1 signaling (Figure 2B). This is in line with early placental development occurring in a low oxygen environment. We confirmed that TE-like CTB are only present in early time points, as we could not detect EPCAM-positive cells in second and third trimester tissue (Supplementary Figure 3). Opposite to cycling CTBs, TE-like CTB show upregulation of *MKI67* but not *PCNA* and are mainly in the G2M phase rather than S phase (Figure 2C). This suggests that TE-like are highly proliferative and actively dividing.

**Figure 2.**
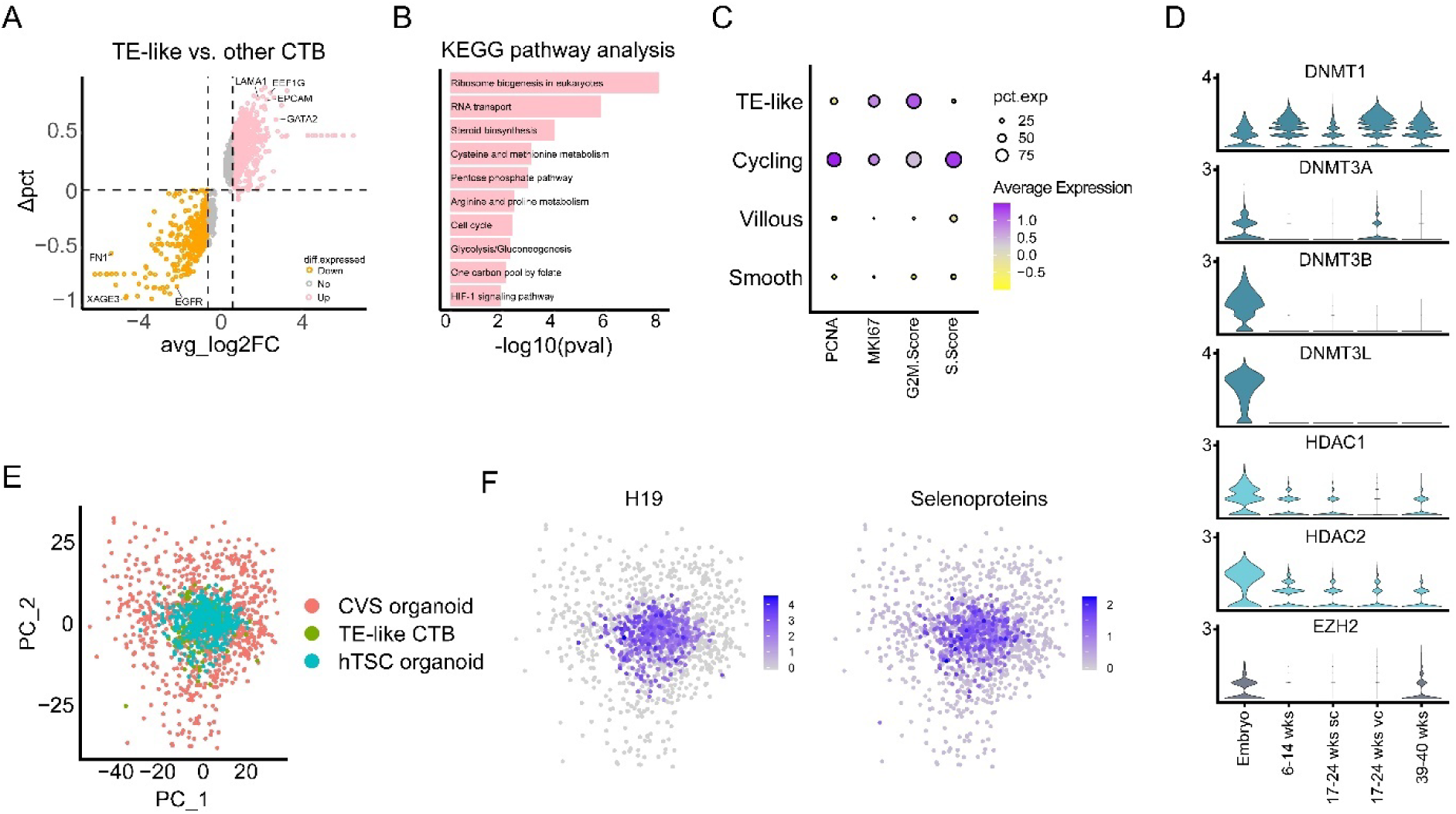
TE-like CTB resemble a transit-amplifying primitive trophoblast population. (A) Vulcano plot showing differentially expressed genes between TE-like CTB and all other CTB clusters. The y-axis displays the difference in the percentage of cells expressing the gene. (B) KEGG pathway enrichment analysis of the differentially expressed genes in TE-like CTB. (C) Dot plot showing average expression of proliferation (*PCNA* and *MKI67*) and cell cycle markers in all CTB clusters. (D) Violin plots showing the average expression of epigenetic regulators for each gestational timepoint. (E) PCA plot showing integration of single-cell transcriptional profiles of TE-like CTB, CVS-derived trophoblast organoids and hTSC-derived trophoblast organoids and (F) expression of H19 and selenoproteins.

Additionally, TE-like CTB showed high expression of de novo methylases *DNMT3A*, *DNMT3B* and *DNMT3L* and epigenetic regulators *HDAC1/2* and *EZH2* (Figure 2D). De novo methylation regulates early placental development and function by establishing the trophoblast methylome, while HDAC1 is known to control lineage-specific transcriptional networks in trophoblast stem cells (TSCs) (18, 19). We compared the transcriptome of TE-like CTB with those of CTB populations from publicly available scRNA-seq datasets of trophoblast organoid cultures, derived from either chorionic villi (CVS organoid) (20) or human TSCs (hTSC organoid) (21). Principal component analysis (PCA) from the integrated CTB transcriptomes showed that TE-like CTB highly correspond with CTBs from hTSC organoids (Figure 2E). Highly expressed transcripts shared between TE-like CTB and hTSC organoids and significantly different to CVS organoids, related amongst others to the H19 complex and the selenoprotein family (Figure 2F). Altogether, this data suggests that TE-like CTB resemble a transit-amplifying primitive trophoblast population.

### Smooth CTB form a columnar epithelium devoid of syncytialization

As the dataset of Marsh and colleagues also sampled the smooth chorion, we could perform a comparison between CTB populations found at distinct locations in the placenta (16). By integrating all datasets, we could use the specific transcriptome of cells from the smooth chorion to identify smooth CTB populations at different gestational time points. Interestingly, all datasets contained smooth CTB cells. Compared to CTB in the villous chorion, smooth CTBs showed upregulation of *SLC2A3*, *CD9*, *EGLN3* and *ITGA2* (Figure 3A). Smooth CTB showed depletion of *BCAM*, *SMAGP* and *DUSP9*, which are all upregulated in villous CTB. Functional enrichment analysis showed that smooth CTB differentially express transcripts related to cell adhesion, tumor biology and HIF-1 signaling (Figure 3B). Additionally, Gene Set Enrichment Analysis (GSEA) confirmed cell adhesion and response to hypoxia as top pathways, with Rap1 signaling and FoxO signaling as effector pathways, respectively (Figure 3C). CellChat analysis indicated paracrine JAM signaling in smooth CTB, with Junctional Adhesion Molecule A (JAM-A/*F11R*) as main contributor (Figure 3D). JAM-A is an important regulator of epithelial paracellular permeability (22). Smooth CTB showed high expression of columnar epithelium markers *KRT8* and *KRT18* and downregulation of syncytiotrophoblast differentiation markers *CGA* and *ERVW-1* (Figure 3E). All these data combined indicate that smooth CTB form a leak-tight epithelial layer that adapts to a low oxygen environment and do not undergo syncytialization. This observation appears to be consistent with the avascular identity of the smooth chorion, as it lacks extensive blood vessel development compared to the highly vascularized villous chorion.

**Figure 3.**
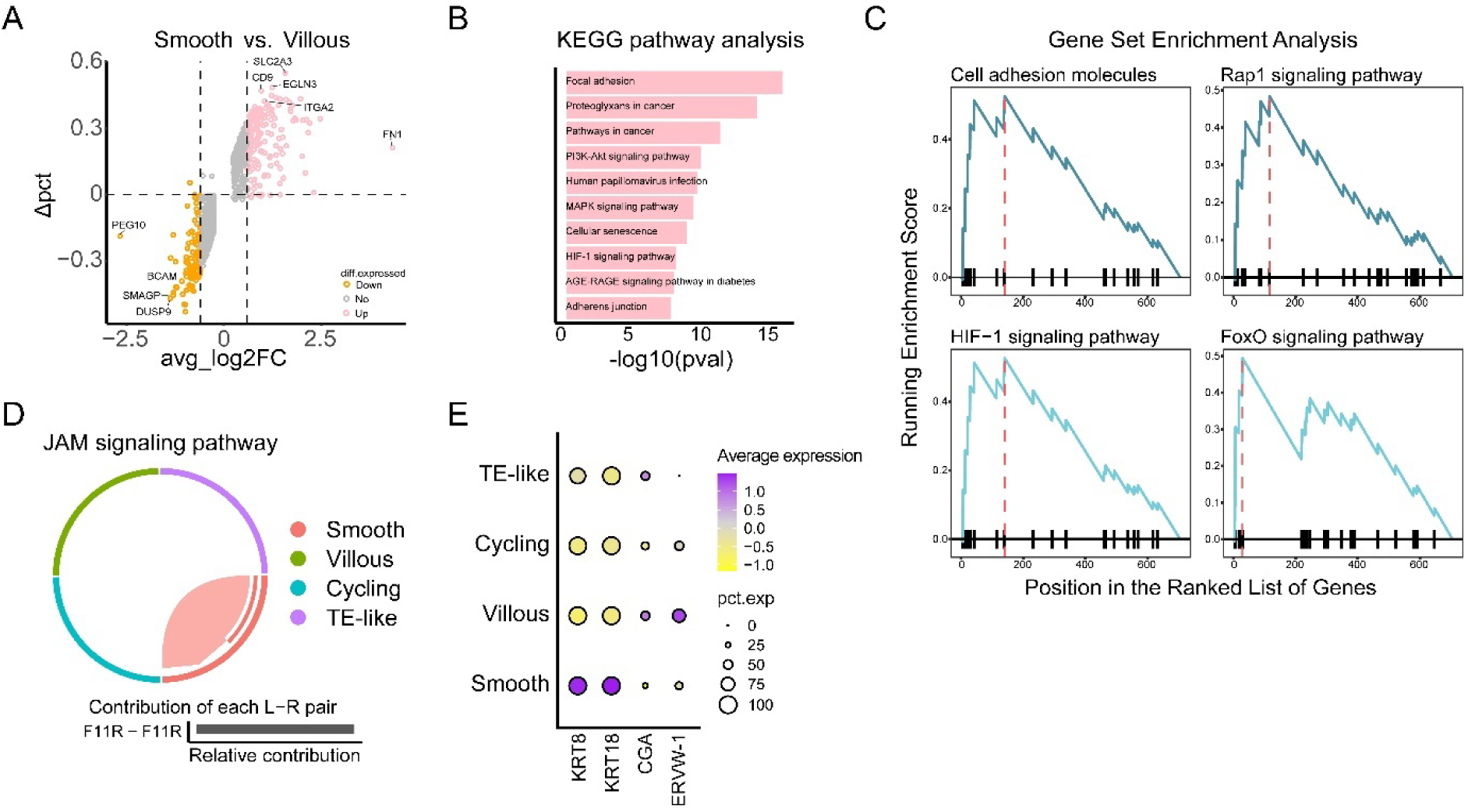
Smooth CTB form a columnar, leak-tight epithelial layer that adapts to a low oxygen environment and does not undergo syncytialization. (A) Vulcano plot showing differentially expressed genes between smooth CTB and villous CTB. The y-axis displays the difference in the percentage of cells expressing the gene. (B) KEGG pathway enrichment analysis of the differentially expressed genes in smooth CTB. (C) Gene Set Enrichment Analysis of genes differentially expressed in smooth CTB. (D) Chord diagram showing autocrine JAM signaling in smooth CTB detected by CellChat analysis. Bar plot indicates the contribution of the involved receptor-ligand pairs. (E) Dot plot showing the average expression of *KRT8* and *KRT18* (columnar epithelial markers) and *CGA* and *ERVW-1* (syncytiotrophoblast differentiation markers) in all CTB clusters.

### BCAM marks CTB cells with progenitor abilities in the villous chorion throughout gestation

To further characterize the villous CTB population, we validated BCAM as a marker for CTB populations in the villous chorion of both second and third trimester placentas. Transcriptionally, *BCAM* was differentially expressed in villous CTB compared to the other CTB populations. Staining of serial sections of placental villi showed that BCAM colocalized with ITGA6, a consensus marker for CTB cells, with BCAM expressed on the basolateral side of the cells (Figure 4A). BCAM expression was consistent in all villi, lining the cell layers underneath the syncytium (marked by CGB) (Figure 4B). To test whether BCAM^+^ CTB in the villous chorion display progenitor capacity, we isolated them from term placentas and initiated organoid cultures. For this, we dissociated placentas to obtain high viability cell suspensions (∼80% viability), followed by magnetic-activated cell sorting to sort out BCAM^+^ CTB (Supplementary Figure 4A-C). Roughly 10% of the cell suspension was positive for BCAM (Supplementary Figure 4B). The sorted BCAM^+^ cells were embedded in Matrigel and overlayed with previously defined trophoblast organoids medium. After roughly 7 days, small organoid structures developed that further grow out as villous-like organoids (Figure 4C). Whole mount staining showed that syncytium (marked by GDF15) is formed in the center of the organoids (Figure 4D), suggesting that BCAM^+^ CTBs could indeed function as trophoblast progenitors with potential for syncytiotrophoblast differentiation. Although we observed the formation of some organoid structures when embedding BCAM^-^ cells, the BCAM^+^ fractions generated a significantly greater number of organoid structures (Supplementary Figure 4D). This indicates that the isolation of BCAM^+^ CTBs might enhance the efficiency of trophoblast organoid derivation from term placentas.

**Figure 4.**
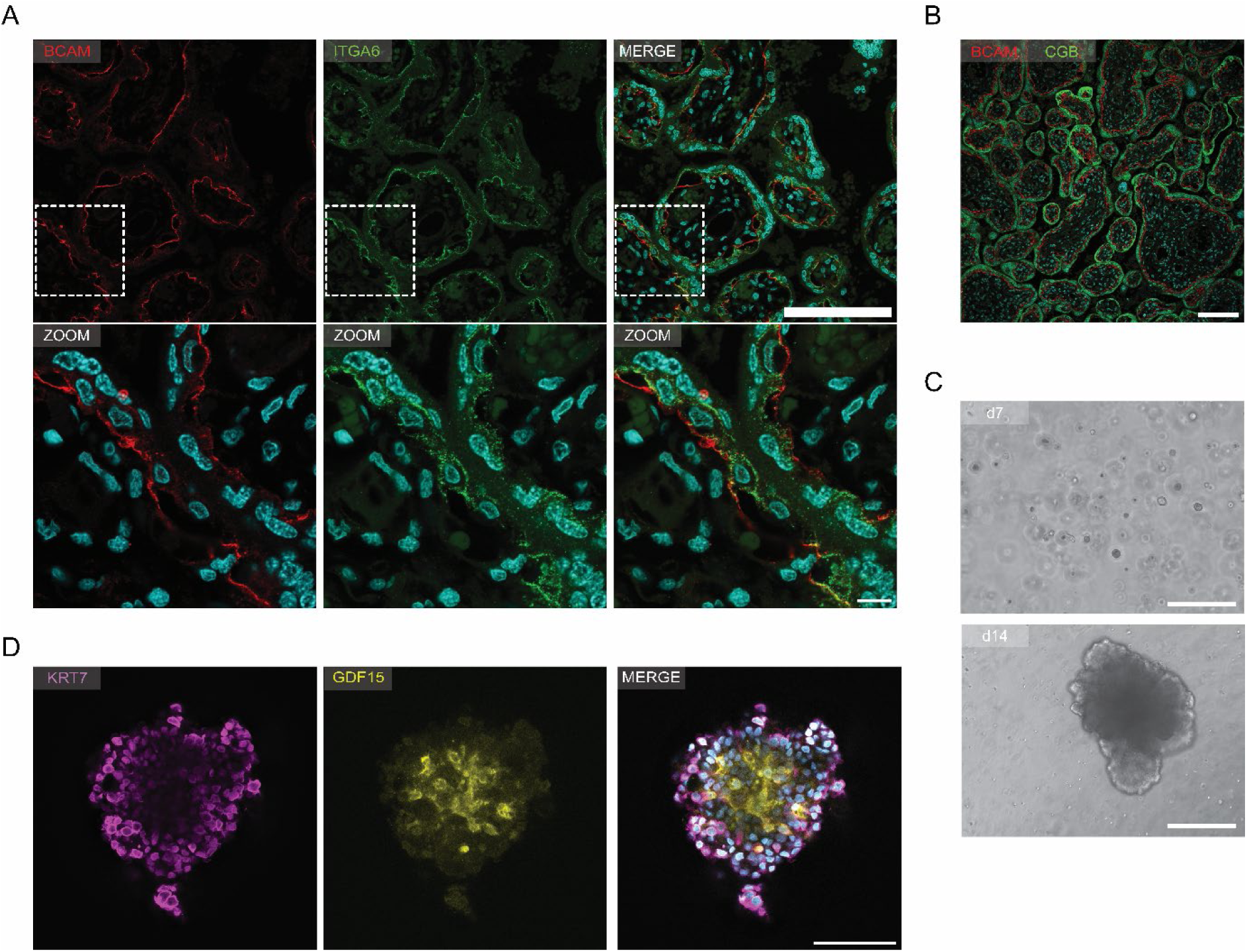
BCAM is a marker for CTBs possessing progenitor capacity within the villous chorion. (A) Immunostaining of BCAM (red) and ITGA6 (green) in term placental tissue. Scale bar, 500 µm and 50 µm (zoom). (B) Immunostaining of BCAM (red) and CGB (green) in term placental tissue. Scale bar, 500 µm. (C) Brightfield images of trophoblast organoid cultures derived from BCAM^+^ cells isolated from term placental tissue at day 7 and day 14. Scale bars, 200 µm. (D) Immunostaining of KRT7 (purple) and GDF15 (yellow) in trophoblast organoids derived from BCAM^+^ cells at day 14. Scale bar, 200 µm.

### BCAM colocalizes with LAMA5 at the basal lamina

The staining pattern of BCAM, together with BCAM (Basal Cell Adhesion Molecule) being a membrane-bound glycoprotein, led us to hypothesize that contact of CTB with the basal lamina in placental villi might play a role in regulating CTB fate (Figure 5A). Electron microscopy confirmed that CTB situated below the syncytium are in contact with a basal membrane (Figure 5B). This basal membrane forms a barrier between the trophoblast layers and stromal core of the villi. Protein prediction analysis using STRING indicated LAMA5 as top interaction partner of BCAM (Figure 5C). Staining of serial sections of placental villi indeed demonstrated colocalization of BCAM and LAMA5 throughout the entire villi (Figure 5D). This suggests that LAMA5 is a key component of the basal lamina in placental villi. To investigate whether trophoblast organoids express BCAM on their periphery, where the cells are in contact with the laminin-rich hydrogel (representing the basal lamina), we quantified the localization of BCAM. Indeed, BCAM is expressed on the outside of the organoids (Figure 5E). Overall, this suggests that BCAM^+^ CTBs are in contact with the basal lamina in placental villi.

**Figure 5.**
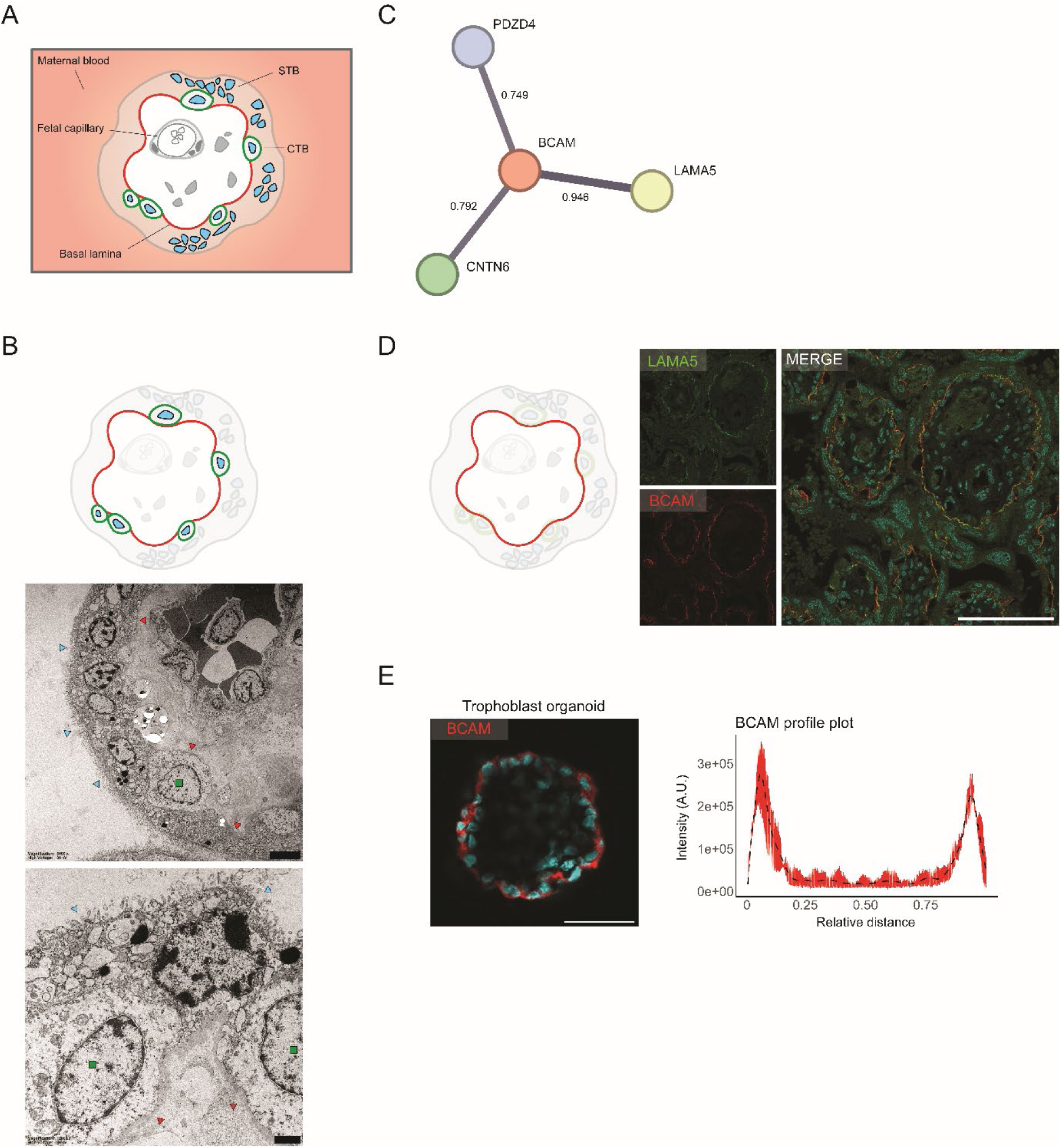
Colocalization of BCAM and LAMA5 suggests contact of progenitor CTBs with basal lamina. (A) Schematic cross section of chorionic villi within the placenta. CTB; cytotrophoblast, STB; syncytiotrophoblast. (B) Electron microscopy images of term placental tissue. Blue arrowheads indicate the syncytium with multiple nuclei, green squares indicate cytotrophoblast, and red arrowheads indicate the basal lamina. Scale bars, 5 µm (top) and 1 µm (bottom). (C) STRING protein network analysis shows top 3 interactors of BCAM with interaction scores. Thickness of lines indicate confidence. (D) Immunostaining of BCAM (red) and LAMA5 (green) in term placental tissue. Scale bar, 500 µm. (E) Immunostaining of BCAM (red) in trophoblast organoids at day 14. Scale bar, 200 µm. Quantification of BCAM intensity profiles measured across multiple line scans (n = 3) per organoid (n = 10). Trend analysis is performed using orthogonal polynomial fitting (dashed line).

### Proteomic profiling of decellularized villi across gestation

To further study the basal lamina and other extracellular matrix (ECM) factors as part of a potential niche for CTB cells in placental villi, we decellularized placental villi from both first-trimester and term placentas. Hematoxylin and eosin staining and DNA quantification before and after the decellularization process confirmed that most of the cells were removed from the placental tissue (Supplementary Figure 5A-C). We determined the protein contents of the decellularized samples by mass spectometry. While *BCAM* expression in villous CTBs gradually decreased as pregnancy advances towards term (Figure 6A), quantitative proteomics analysis showed that BCAM protein levels were significantly elevated in decellularized villi of term placentas compared to those from the first trimester (Figure 6B). A similar trend was observed for LAMA5, suggesting that both proteins remain abundant in placental villi throughout the entire pregnancy (Figure 6B). Furthermore, we inferred the potential composition of the basal lamina in decellularized villi of term placentas based on the significant abundance of common basal lamina-associated proteins (23, 24). These included various collagens, including COL6A1-3, COL14A2 and COL4A2; laminins, including LAMA2, LAMC1, LAMB1-2 and LAMA5; as well as nidogen (NID1), fibronectin (FN1), and perlecan (HSPG2) (Figure 6C).

**Figure 6.**
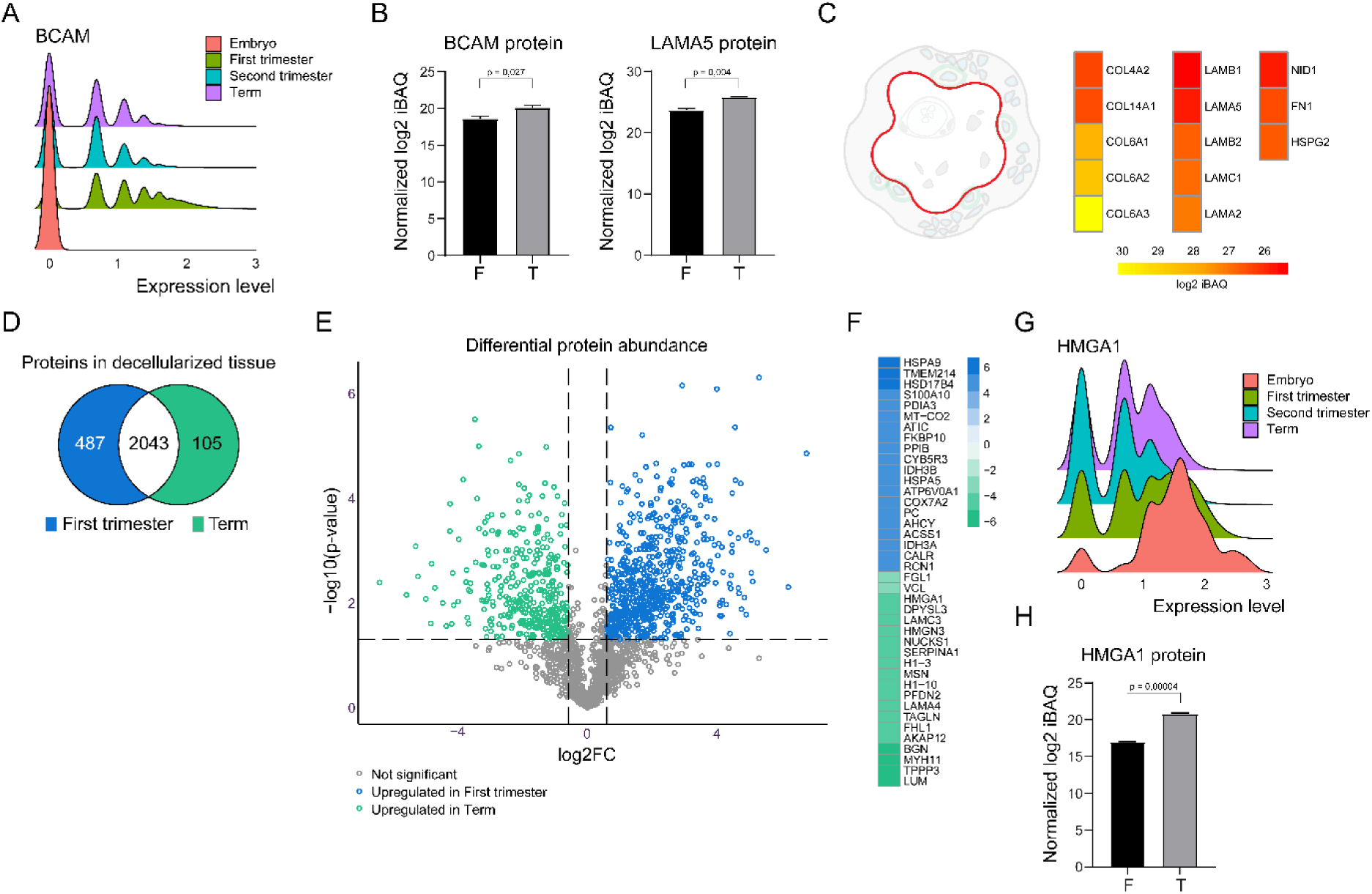
Proteomic profiling of decellularized villi across gestation. (A) Ridgeline plot showing average *BCAM* expression in villous CTB cells throughout gestation. (B) Bar plots showing protein abundance of BCAM and LAMA5 in decellularized villi from first trimester (n = 3) and term (n = 3) placentas. P-values are calculated using a Welch’s t-test. (C) Abundance of basal lamina-associated proteins in decellularized villi from term placentas (n = 3). (D) Venn diagram showing the number of unique or shared proteins in decellularized villi from first trimester and term placentas. Proteins present in all samples (n = 3 each) at either or both gestational times were retained. (E) Vulcano plot showing differential protein abundance of the 2043 proteins shared between decellularized villi from first and term placentas. (F) Fold change of the top 20 differentially abundant proteins in decellularized villi from either first trimester or term placentas. (G) Ridgeline plot showing average *HMGA1* expression in villous CTB cells throughout gestation. (H) Bar plot showing protein abundance of HMGA1 in decellularized villi from first trimester (n = 3) and term (n = 3) placentas. P-value is calculated using a Welch’s t-test.

To explore gestation-related differences in ECM composition and potential niche-related factors, we profiled the complete set of proteins identified across all samples of first trimester and/or term decellularized villi. A majority of the proteins identified (2043; 77.5%) overlapped between both gestational timepoints (Figure 6D). We identified 487 and 105 unique proteins in first trimester and term decellularized villi, respectively. Functional enrichment analysis revealed that the unique proteins identified only in first trimester samples are mostly involved in metabolic processes and neurodegenerative diseases (Supplementary Figure 6A), whereas those identified only in term samples are linked to TNF signaling and atherosclerosis (Supplementary Figure 6B).

Next, we assessed the differences between proteins that were shared between decellularized villi of both gestational time points. Of the 2043 shared proteins, 674 were differentially abundant (≥2-fold, p < 0.05) in first trimester samples, compared to 357 protein in term samples (Figure 6E). Among the top 20 differentially abundant proteins, several classic ECM proteins were more abundant in term samples, including lumican (LUM), biglycan (BGN), and laminins LAMA4 and LAMC3 (Figure 6F). Interestingly, HMGA1, a known regulator of stemness and self-renewal (25, 26), was one of the most differentially abundant proteins at term (Figure 6F). While *HMGA1* expression is highest in embryonic CTB (Figure 6G), it may be secreted, as protein levels are significantly enhanced in decellularized villi at term (Figure 6H). The substantial levels of HMGA1 protein observed in decellularized villi from term placentas suggest an active role for HMGA1 in maintaining the CTB progenitor niche during the later stages of pregnancy.

## Discussion

Previous studies have used single-cell RNA-sequencing to study trophoblast biology in the human placenta (14–17). While these studies have primarily focused on specific trimesters, we aimed to profile CTB dynamics throughout the full gestational period. For this, we pooled these data with embryonic datasets containing early trophoblast cells to systematically characterize the transcriptional landscape of CTB populations throughout the full course of pregnancy.

Our analysis revealed robust transcripts for distinct CTB populations, including a transient stem-cell like subset, CTBs located at the smooth chorion, and BCAM^+^ CTB in the villous region. Notably, this last subset could be isolated from term placental tissue and displayed progenitor abilities in organoid cultures. Based on the localization and pattern of BCAM expression, we hypothesized that BCAM^+^ CTBs maintain a progenitor role late into pregnancy, potentially regulated by their interaction with the underlying basal lamina. This finding supports the notion that stem cell niches, defined by their extracellular environment and signaling cues (27), might continue to influence CTB fate in the placenta up to term. Although stem cell niches have been well characterized in various tissues (28–30), the niche for CTBs remains largely understudied, particularly during later stages of pregnancy.

Previous studies have debated that the number of proliferative CTBs decreases in the placenta over time, with the steepest decline occurring at the end of the first trimester (9, 10). This aligns with the rapid expansion of the placenta in this period, where rapid proliferation of trophoblasts is crucial for securing the conceptus within the uterine wall and forming a robust cytotrophoblastic shell (31). We observed a similar pattern in our analysis, with cycling activity decreasing throughout gestation, but with proliferative CTBs persisting up to term. In the embryonic datasets, we identified a population of *EPCAM*-positive highly proliferative CTBs. EPCAM is a well-known marker for stem and progenitor cells, including in placental tissue (32, 33). Interestingly, we did not detect any EPCAM-positive cells in second-trimester or term placental tissues. Recent analysis of trophoblast populations also restricted *EPCAM* expression to hTSC-derived states (34). Altogether, this may suggest that EPCAM indicates a transit-amplifying population of CTB cells, which gradually lose *EPCAM* expression as they mature. Similar dynamics are observed in other tissues; *EPCAM* is highly expressed in human hepatic stem and progenitor cells but absent in mature hepatocytes (35–37).

Our study highlights the potential involvement of BCAM in regulating CTB progenitor fate throughout gestation. BCAM has previously been implicated as a marker of a primitive trophoblast state in early placental tissue (15). Interestingly, the identification of BCAM as a marker of progenitor cells is not limited to CTBs, as similar roles for BCAM have been observed in other epithelial tissues. For instance, *BCAM* expression has been linked to progenitor or stem cell populations in cornea, lung and thymus (38–40). As an adhesion molecule with a strong interaction with laminin subunit α5, BCAM parallels broader observations of stem cells across tissues being characterized by specific integrins, where integrin expression influences cell proliferation and differentiation by anchoring these cells within a niche (41, 42). Similarly, we hypothesize that the BCAM-laminin interaction at the basal lamina likely plays a role in maintaining the CTB progenitor pool within the placenta, ensuring proper placental development and function throughout pregnancy.

The relevance of these findings extends to pregnancy complications such as preeclampsia and fetal growth restriction, which are associated with placental abnormalities like distal villous hypoplasia and accelerated villous maturation (43, 44). In these conditions, CTB progenitor pools may become depleted early, resulting in aberrant villous development and loss of trophoblast turnover (45). Notably, a recent study implicated that BCAM expression is decreased in preeclamptic placentas, with BCAM deficiency inducing a preeclampsia-like phenotype in rats (46). Future research is needed to determine whether BCAM indeed plays a role in supporting the CTB progenitor niche, potentially by mediating specific cell-ECM interactions, especially in the context of placental abnormalities. Better understanding CTB progenitor niche factors could also offer new strategies to address conditions linked to placental insufficiency. Emerging research has demonstrated that injecting niche factors targeting the ECM and specific receptors can modulate tissue regeneration (47). For instance, enhancement of β1-integrin activity using monoclonal antibodies or treatment with fetal laminin-111 improves muscle regeneration (48, 49). These regenerative strategies could be adapted to support placental regeneration by restoring the CTB progenitor niche, improving trophoblast proliferation and overall placental function.

## Materials and Methods Tissue collection

The collection and utilization of tissues adhered to the regulatory framework outlined by the Erasmus University Medical Center Ethics Board (study number: OZBS71.19172) and were conducted in accordance with the guidelines set forth in the Declaration of Helsinki 2000. Informed consent was obtained from each patient donating placental tissue.

## Bioinformatic analysis

Raw sequencing data were retrieved from the NCBI Geo repository (see Data Availability section). For quality control, cells with fewer than 500 counts, 250 detected genes and low complexity (log10 value of number of genes detected per count < 0.8) were removed. Genes with zero expression in all cells and that are expressed in less than 10 cells were removed. Downstream data analyses were performed using the R package Seurat (version 5.0.1; (50)). Datasets were normalized by the ‘SCTransform’ function. Cell cycle heterogeneity was scored and differences between S and G2M cell cycle phases were regressed out using the functions ‘CellCycleScoring’ and ‘ScaleData’. Integration of the datasets was performed according to the Seurat alignment workflow using the Leiden algorithm. UMAP analysis was performed using the ‘RunUMAP’ function including 20 principal components. Clusters were identified using the ‘FindClusters’ function with a resolution of 0.1. Cluster marker genes were identified by differential expression analysis using the ‘FindAllMarkers’ function with a minimum log fold change value > 0.25, adjusted *p*-value < 0.01 and genes detected in a minimum of 25% cells in each cluster. Gene overrepresentation (GO) analysis on differentially expressed genes was performed using the ‘DEenrichRPlot’ function and the KEGG database. Gene set enrichment analysis (GSEA) on differentially expressed genes was performed using the clusterProfiler package (51) and the KEGG database. Analysis of signaling pattern was performed using the CellChat package with default parameters (52).

## Immunohistochemistry and confocal microscopy

Immunofluorescent staining was performed on formalin-fixed paraffin-embedded placental tissue sections (18–39 weeks of gestation). Tissue slides were dried overnight at 37°C, washed 3×5 min in xylene, 3×5 min in 100% ethanol and 3×5 min in PBS. Antigen retrieval was performed by heating the slides in a microwave for 20 min in 10 mM sodium citrate buffer (pH 6.0, 0.05% Tween). Sections were blocked in 10% donkey serum/5% BSA/PBS for 30 min at room temperature. Primary antibodies were diluted in 1% BSA/PBS and incubated overnight at 4°C. The following primary antibodies were used: polyclonal anti-BCAM (rabbit, 1:200, Sigma, HPA005654), monoclonal anti-ITGA6 (mouse, 1:100, Sigma, AMAB91450), monoclonal anti-hCG (mouse, 1:200, Abcam, AB9582), and monoclonal anti- LAMA5 (mouse, 1:200, Sigma, SAB4501720). Fluorescently-labeled secondary antibodies (Alexa Fluor, 1:500, Invitrogen, A32733) were diluted in BSA/PBS and incubated for 1 h at room temperature. Autofluorescence was quenched using the Vector trueVIEW (Vector Laboratories) kit according to the manufacturer’s instructions. All slides were mounted using ProLong Gold mounting medium containing DAPI (Life Technologies) and images on a Leica Stellaris 5 KEA confocal microscope. All images were acquired using a Plan-Apochromat 20x objective with 0.8 NA or a Plan-Neofluar 40x oil immersion objective with 1.3 NA. Images were processed with ImageJ.

## Isolation of BCAM^+^ cells from placental tissue

Placental tissue (37-40 weeks of gestation) was dissociated to obtain single cell suspensions as previously described (53). Briefly, 2-3 grams of tissue was cut from the basal plate and small pieces of 1 mm^2^ were scraped carefully using a scalpel and collected in gentleMACS C tubes (Miltenyi Biotech). Enzymes from the Umbilical Cord Dissociation kit (Miltenyi Biotech) were added according to the manufacturer’s instructions and tubes were incubated at 37°C for 3 h. Tubes were gently inverted every 30 min to ensure thorough digestion of the sample. After incubation, tubes were placed on a gentleMACS Octo Dissociator (Miltenyi Biotech) and mechanically dissociated using program h_cord_01. The obtained suspensions were filtered through a 100 µm cell strainer and remaining red blood cells were removed using ACK lysing buffer. Cells were counted using Trypan Blue to assess viability. Cell suspensions were either directly using for magnetic-activated cell sorting (MACS) or cryopreserved in Cellbanker-2 (Amsbio). To isolate BCAM^+^ cells using MACS, cells suspensions were centrifuged at 300 x *g* for 5 min and resuspended in 100 µl ice-cold MACS buffer (1% BSA, 2 mM EDTA in PBS). Cell suspensions were incubated with 5 µL of anti-BCAM PE-conjugated antibody (Miltenyi Biotech, 130-126-539) in the dark for 10 min at 4°C. Cell suspensions were washed twice with MACS buffer, centrifuged at 300 x *g* for 5 min and resuspended in 90 µl ice-cold MACS buffer. Next, cell suspensions were incubated in 10 µL of PE microbeads (Miltenyi Biotech) for 15 min at 4°C.

Cell suspensions were washed with MACS buffer, centrifuged at 300 x *g* for 5 min and resuspended in 500 µl ice-cold MACS buffer. Cell suspensions were then applied onto LS Columns (Miltenyi Biotech) stationed in a MidiMACS magnet (Miltenyi Biotech) according to the manufacturer’s instructions. Both BCAM^+^ and BCAM^-^ fractions were collected, counted and used for organoid cultures.

## Organoid cultures

Cell suspensions were filtered through a 40 µm cell strainer, washed in Advanced DMEM/F12 medium (Gibco) and centrifuged at 300× *g* for 5 min. Cells were plated in GFR Matrigel (Corning) and cultured in trophoblast organoid medium (TOM) consisting of Advanced DMEM/F12 supplemented with 1X B27 minus vitamin A (Life Technologies), 1X N2 (Life Technologies), 10% Knockout serum replacement (vol/vol), 100 µg/mL primocin (Invivogen), 2 mM L-glutamine (Sigma), 1.25 mM N-Acetyl-L-cysteine (Sigma), 10 mM nicotinamide (Sigma), 500 nM A83-01 (Tocris), 1.5 µM CHIR99021 (Tocris), 50 ng/mL recombinant human EGF (Gibco), 80 ng/mL recombinant human R- spondin1 (Peprotech, London, UK), 100 ng/mL recombinant human FGF2 (Peprotech), 50 ng/mL recombinant human HGF (Gibco), 5 µM Y--27632 (Merck Millipore), and 2.5 µM prostaglandin E2 (Sigma), as previously described (54). Medium was replaced every 2–3 days and organoids were passaged every 7 to 10 days depending on their size and density. Whole-mount immunofluorescence staining of trophoblast organoids was performed as previously described (55). The following primary antibodies were used: monoclonal anti-KRT7 (mouse, 1:100, DAKO, M7018), polyclonal anti-GDF15 (rabbit, 1:100, Sigma, HPA011191), and polyclonal anti-BCAM (rabbit, 1:200, Sigma, HPA005654). Before imaging, organoids were embedded in Matrigel drops in the center of 60 mm Petri dishes. Imaging was performed using the Leica SP5 Intravital confocal microscope with a HCX APO 20x water dipping objective. Further analysis was performed using ImageJ software.

## Decellularization of placental tissue

First trimester and term placental tissues were collected and stored at -20°C until start of the decellularization experiment. Upon collection, additional biopsies (∼1-2 cm³) were snap frozen or stored in PFA for either DNA analysis and histology, respectively. Placental tissue was thawed at 4°C for at least 16 hours. Next, tissue was cut in pieces of 1-2 cm² and roughly 10-15 g of tissue was used per experiment. Excessive blood was removed by washing the tissue in water, followed by 2×30 min in PBS. For decellularization, tissue was washed in hypertonic saline (9% NaCl) for 1 h, in PBS for 30 min, and 10×1 h in PBS with 4% Triton X-100 and 1% NH3 at room temperature. After 3 cycles in, the placental tissue should start to turn white/translucent. After at least 5 cycles, one of the washes with PBS supplemented with 4% Triton X-100 and 1% NH3 was performed overnight. After 10 cycles, the tissue is washed in PBS for 1 h, followed by 3.5 h in DNAse buffer (0.9% NaCl, 100 mM CaCl2, 100 mM MgCl2 and 2 mg DNAse) at 37°C. Finally, the tissue is washed twice in PBS for 30 min at room temperature. All washes were performed in a glass spinner flask under constant agitation (1000 RPM) using a minimal volume of 250 mL. After decellularization, tissues were lyophilized to remove all residual water and stored at -20°C. For analysis, DNA was isolated using the PicoPure™ DNA extraction kit (Thermo Scientific) following the manufacturer’s instructions and concentrations were measured on a Qubit™ Fluorometer (Thermo Scientific). Samples with a DNA concentration lower than 20 ng dsDNA/mg wet tissue were considered sufficiently decellularized. For histology, samples were fixed in PFA and stained with hematoxylin and eosin.

## Mass spectrometry

Lyophilized decellularized tissue was prepared for mass spectrometry using a modified SPEED protocol (56). Sample masses were determined roughly and ranged from approximately 0.9 to 2.5 mg. The lyophilized powders were transferred to 1.5 mL Bioruptor® Pico vials (Diagenode) and sample tubes were rinsed with a volume of 100% TFA where the volume (μl) used was calculated as 80% x mass (mg) x 160 μl/mg. The vials were capped, sealed with Parafilm, and incubated at 40°C and 300 rpm on a thermomixer for 4 days until clear. Samples were sonicated in a Bioruptor Pico® (Diagenode) for 10 minutes at 30 sec on/off intervals daily. TFA was supplemented if necessary and vials were replaced when integrity was affected. An aliquot of 20 μl of solubilized sample (estimated at 120 µg protein) was supplemented with 2M Tris base until the pH was between 7.5 and 9, followed by the addition of 10 μl of 1M NaCl, and the final volume was adjusted to 200 μl with water. Next, proteins were precipitated according to the SP4 method (57). To the protein aggregates, 100 μl of 0.5% sodium deoxycholate (SDC), 50mM triethylammonium bicarbonate (TEAB) and 2.5 µg LysC (Fujifilm Wako Chemicals) were added and the samples were incubated at 37°C at 1100 rpm for 2 hours. Subsequently, 20 μl of immobilized trypsin (TPCK-treated, Thermo Scientific) was added, and the incubation was continued overnight at 30°C with shaking at 1100 rpm. The following day, 10% TFA was added until the pH dropped below 3 to precipitate the SDC. After mixing and centrifugation, the supernatants were collected. Approximately 50 μg of protein was removed and adjusted to a final volume of 100 μl with 0.1% TFA. The peptides were then cleaned using a homemade double-plugged Empore™ C18 StageTip (3M) and subsequently dried. The dried peptides were redissolved in 0.5% formic acid and 2% acetonitrile, containing 0.5 iRT peptides (Biognosys), and analyzed by nanoflow liquid chromatography-tandem mass spectrometry (nLC-MS/MS) on an EASY-nLC system coupled to an Orbitrap Fusion™ Lumos™ Tribrid™ Mass Spectrometer (Thermo Scientific), as previously described (58). Test injections were performed using a DDA method and analyzed with MaxQuant (v.2.4.2.0) using standard settings. Peptide intensities were summed after correcting for contaminants, and injection volumes were normalized to the median summed intensity. All samples were then reinjected using a DIA method. All data was normalized and analyzed using Spectronaut (v.18). Protein groups were inferred using the IDPicker algorithm. P-values were calculated using a Welch’s t-test.

## Acknowledgements

We are thankful to the Department of Clinical Genetics for their help in collecting first-trimester placental tissue. We thank the Department of Pathology for the electron microscopy images of term placental tissue.

---

**Figure S1:**
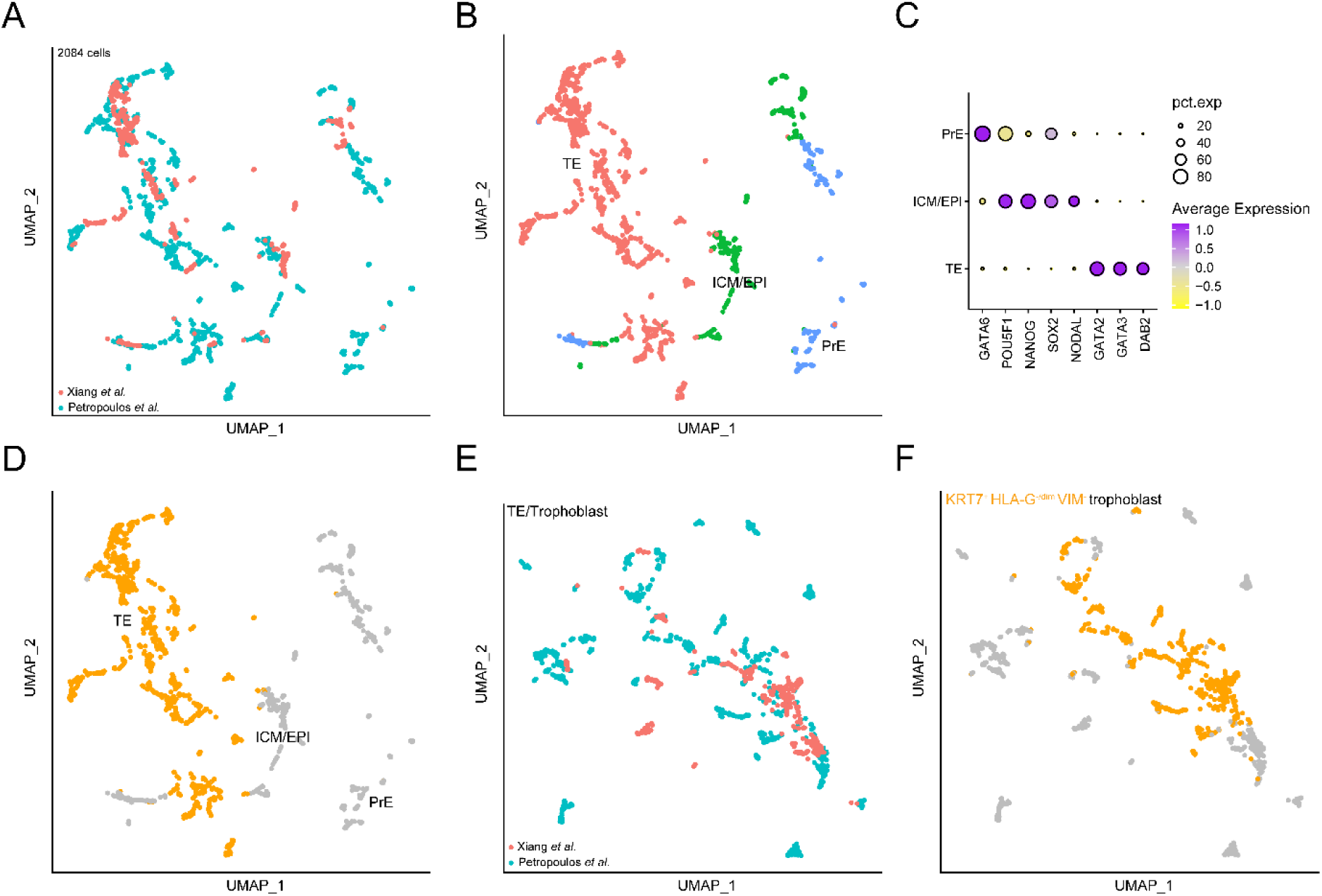
Segregation of trophectoderm lineage from embryonic datasets. (A) UMAP plot showing integration of the human pre-implantation and pre-gastrulation embryo (E3-E12/13) datasets from Petropoulos *et al.* (12) and Xiang *et al.* (13). (B) UMAP plot showing the clustering of trophectoderm (TE; in red), inner cell mass/epiblast (ICM/EPI; in green) and primitive endoderm (PrE; in blue) lineages after integration. (C) Dotplot showing the average expression of markers for the TE, ICM/EPI and PrE lineages. (D) UMAP plot showing the selected TE population (in orange) that was segregated using TE markers *GATA2*, *GATA3* and *DAB2*. (E) UMAP plot showing the re-clustering of the segregated TE population. (F) UMAP plot showing the trophoblast population (in orange) that was segregated from the TE population for further analysis. Trophoblast markers *KRT7*, *ITGA6* and *TEAD4* were used to enrich for cytotrophoblast cells.

**Figure S2:**
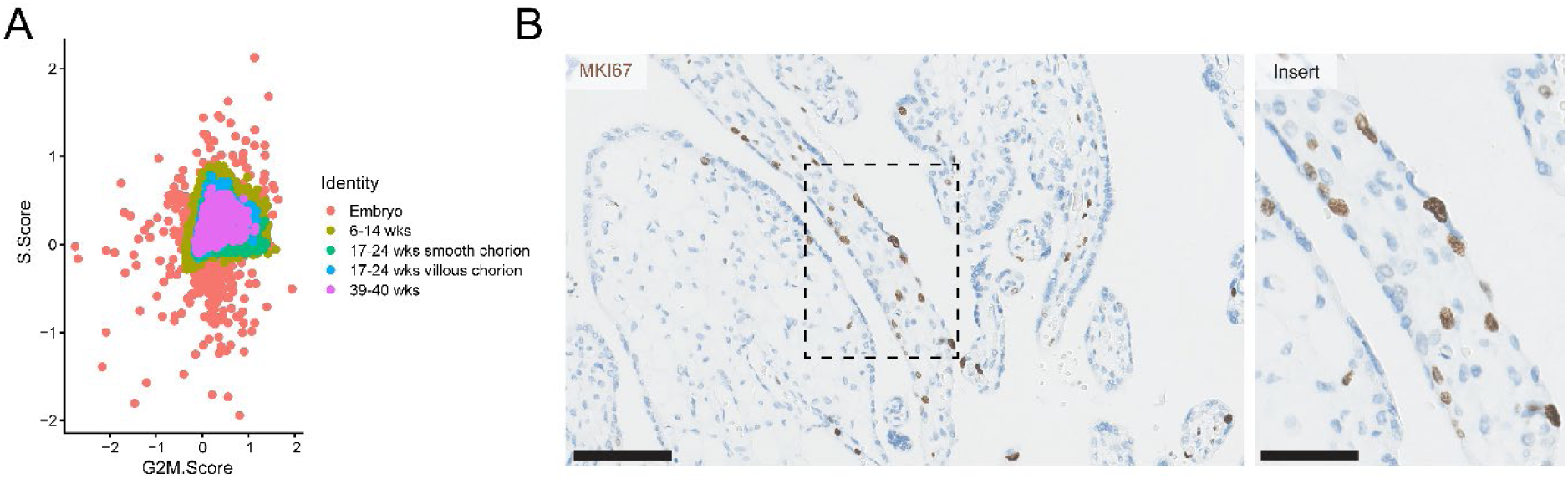
Cell cycle analysis of CTBs during gestation. (A) Scatterplot showing S.Score and G2M.Score (average expression of genes involved in S-phase and G2M-phase of the cell cyle). (B) Immunohistochemical staining for MKI67 (brown) in term placental tissue. Scale bars, 200 µm (left) and 100 µm (insert).

**Figure S3:**
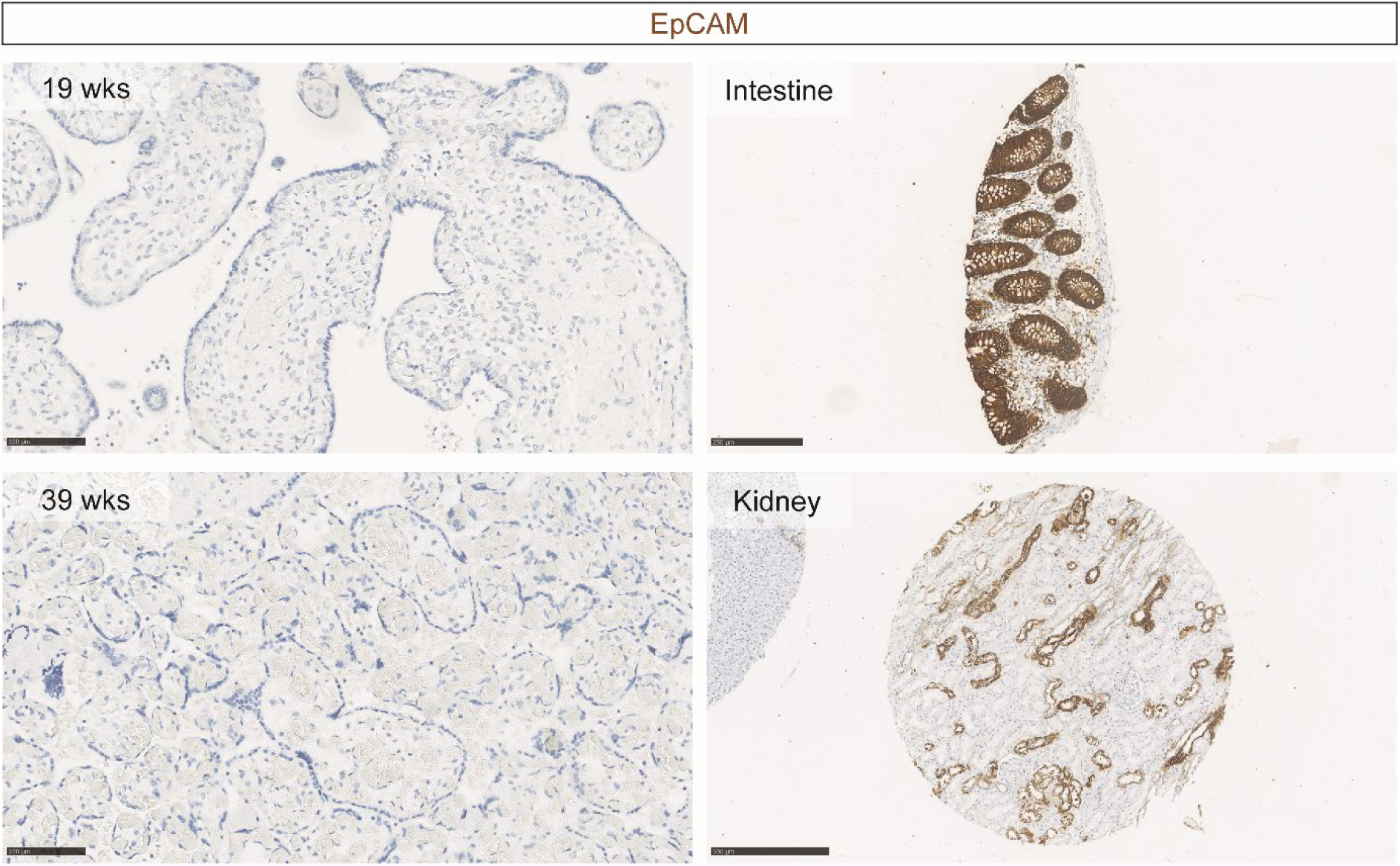
EPCAM staining in placental tissue. Immunohistochemical staining for EPCAM (brown) in second (19 weeks) and term (39 weeks) placental tissue. Intestinal and kidney tissues are used as positive controls. Scale bars are indicated.

**Figure S4:**
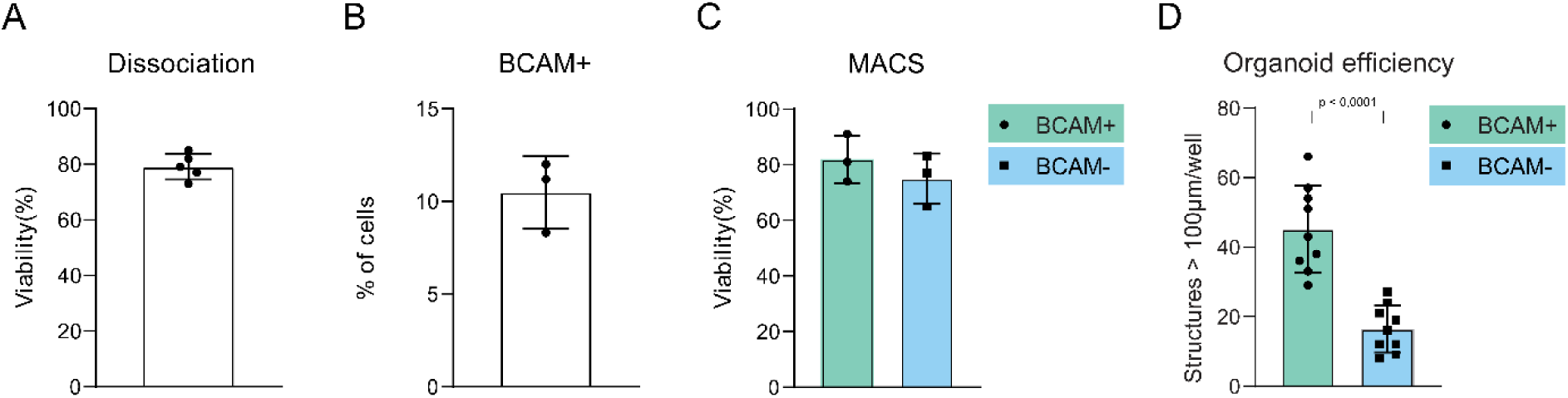
Isolation of BCAM^+^ cells from term placental tissue. (A) Bar plot showing the viability of cell suspensions after dissociation of term placental tissue (n = 5). (B) Bar plot showing the relative number of BCAM^+^ cells isolated using magnetic-activated cell sorting (MACS) from placental cell suspensions (n = 3). (C) Bar plot showing the viability of BCAM^+^ and BCAM^-^ cells after MACS isolation from placental cell suspensions (n = 3). (D) Bar plot showing the number of trophoblast organoid structures larger than 100 µm per well (n = 3 wells per experiment, n = 3 experiments) after day 10 of culture for BCAM^+^ and BCAM^-^ cells. The p-value is calculated using the Mann-Whitney U test.

**Figure S5:**
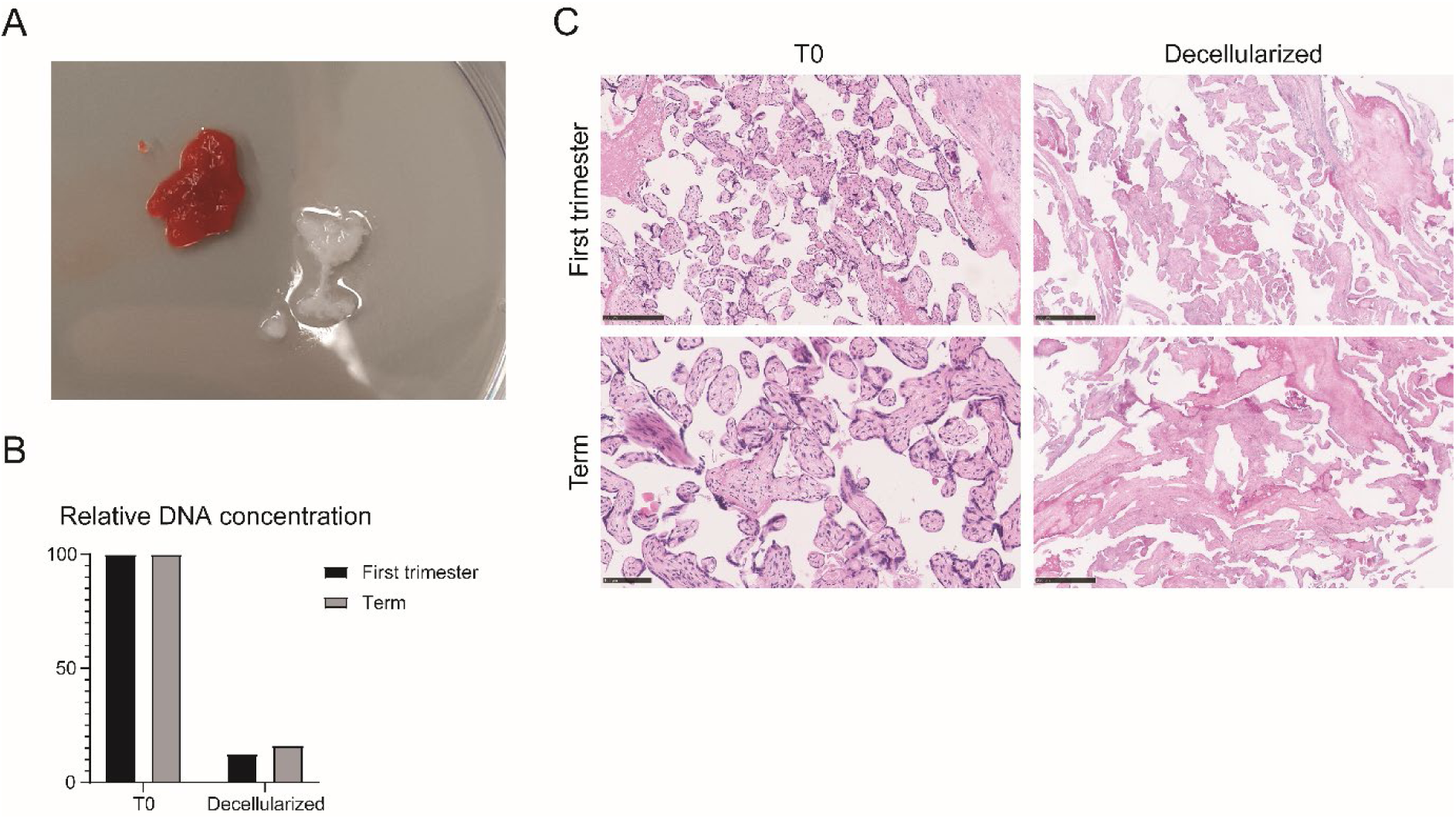
Decellularization of placental tissue. (A) Picture of placental tissue before (red-colored tissue) and after (translucent-colored tissue) decellularization. The resultant decellularized tissue becomes visibly translucent. (B) Bar plot showing relative DNA concentration before (T0) and after decellularization of first trimester (n = 3) and term (n = 3) placental tissue. (C) H&E staining of placental tissue before and after decellularization. Scale bars are indicated.

**Figure S6:**
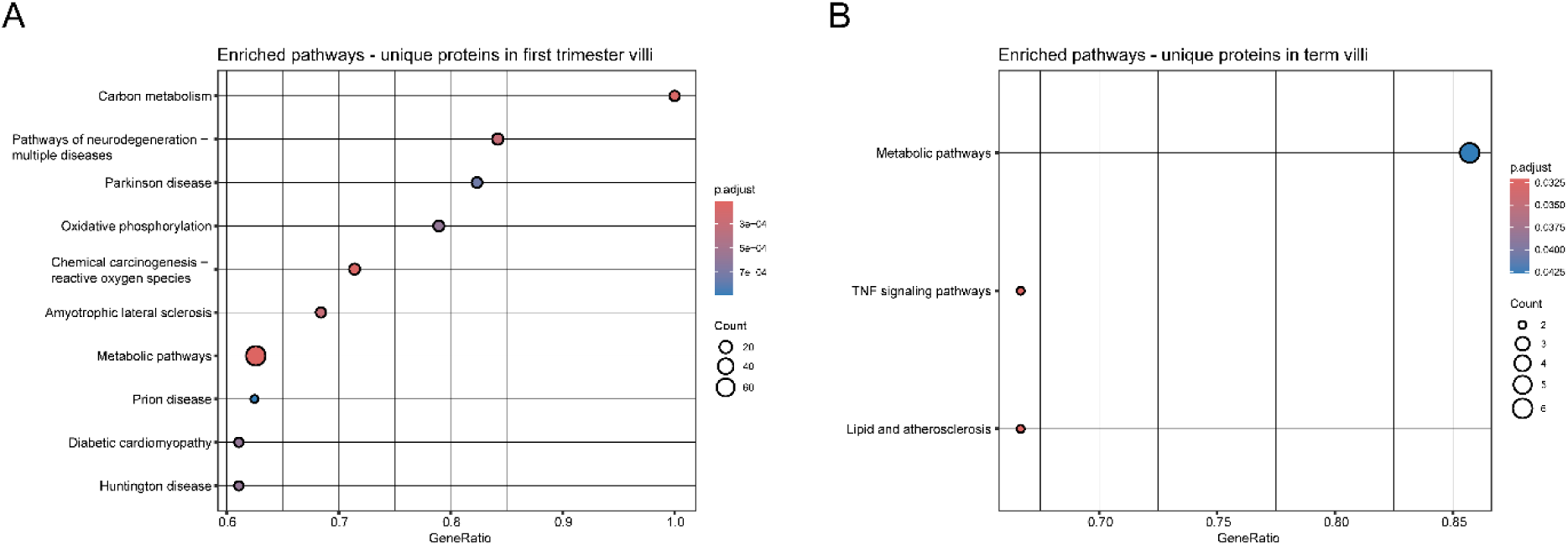
Functional enrichment analysis of unique proteins in decellularized villi. (A) KEGG pathway enrichment analysis of the unique proteins (n = 487) in decellularized villi of first trimester placentas. (B) KEGG pathway enrichment analysis of the unique proteins (n = 105) in decellularized villi of term placentas.

## References

1. Burton GJ, Fowden AL. The placenta: a multifaceted, transient organ. Philos Trans R Soc Lond B Biol Sci. 2015;370(1663):20140066.

2. Aplin JD, Lewis RM, Jones CJP. Development of the Human Placental Villus. Reference Module in Biomedical Sciences: Elsevier; 2018.

3. Turco MY, Moffett A. Development of the human placenta. Development. 2019;146(22).

4. Aplin JD, Myers JE, Timms K, Westwood M. Tracking placental development in health and disease. Nature Reviews Endocrinology. 2020;16(9):479–94.

5. Knöfler M, Haider S, Saleh L, Pollheimer J, Gamage T, James J. Human placenta and trophoblast development: key molecular mechanisms and model systems. Cell Mol Life Sci. 2019;76(18):3479–96.

6. Cross JC. Adaptability and potential for treatment of placental functions to improve embryonic development and postnatal health. Reprod Fertil Dev. 2016;28(1-2):75–82.

7. Sferruzzi-Perri AN, Lopez-Tello J, Salazar-Petres E. Placental adaptations supporting fetal growth during normal and adverse gestational environments. Experimental Physiology. 2023;108(3):371–97.

8. Mayhew TM. Turnover of human villous trophoblast in normal pregnancy: what do we know and what do we need to know? Placenta. 2014;35(4):229–40.

9. Hemberger M, Udayashankar R, Tesar P, Moore H, Burton GJ. ELF5-enforced transcriptional networks define an epigenetically regulated trophoblast stem cell compartment in the human placenta. Human Molecular Genetics. 2010;19(12):2456–67.

10. Burton GJ, Cindrova-Davies T, Turco MY. Review: Histotrophic nutrition and the placental- endometrial dialogue during human early pregnancy. Placenta. 2020;102:21–6.

11. Yang L, Semmes EC, Ovies C, Megli C, Permar S, Gilner JB, et al. Innate immune signaling in trophoblast and decidua organoids defines differential antiviral defenses at the maternal-fetal interface. Elife. 2022;11.

12. Petropoulos S, Edsgärd D, Reinius B, Deng Q, Panula SP, Codeluppi S, et al. Single-Cell RNA-Seq Reveals Lineage and X Chromosome Dynamics in Human Preimplantation Embryos. Cell. 2016;165(4):1012–26.

13. Xiang L, Yin Y, Zheng Y, Ma Y, Li Y, Zhao Z, et al. A developmental landscape of 3D- cultured human pre-gastrulation embryos. Nature. 2020;577(7791):537-42.

14. Vento-Tormo R, Efremova M, Botting RA, Turco MY, Vento-Tormo M, Meyer KB, et al. Single-cell reconstruction of the early maternal–fetal interface in humans. Nature. 2018;563(7731):347-53.

15. Shannon MJ, Baltayeva J, Castellana B, Wächter J, McNeill GL, Yoon JS, et al. Cell trajectory modeling identifies a primitive trophoblast state defined by BCAM enrichment. Development. 2022;149(1).

16. Marsh B, Zhou Y, Kapidzic M, Fisher S, Blelloch R. Regionally distinct trophoblast regulate barrier function and invasion in the human placenta. Elife. 2022;11.

17. Yang J, Gong L, Liu Q, Zhao H, Wang Z, Li X, et al. Single-cell RNA-seq reveals developmental deficiencies in both the placentation and the decidualization in women with late-onset preeclampsia. Front Immunol. 2023;14:1142273.

18. Kidder BL, Palmer S. HDAC1 regulates pluripotency and lineage specific transcriptional networks in embryonic and trophoblast stem cells. Nucleic Acids Res. 2012;40(7):2925–39.

19. Andrews S, Krueger C, Mellado-Lopez M, Hemberger M, Dean W, Perez-Garcia V, et al. Mechanisms and function of de novo DNA methylation in placental development reveals an essential role for DNMT3B. Nature Communications. 2023;14(1):371.

20. Schäffers OJM, Dupont C, Bindels EM, Van Opstal D, Dekkers DHW, Demmers JAA, et al. Single-Cell Atlas of Patient-Derived Trophoblast Organoids in Ongoing Pregnancies. Organoids. 2022;1(2):106–15.

21. Karvas RM, Khan SA, Verma S, Yin Y, Kulkarni D, Dong C, et al. Stem-cell-derived trophoblast organoids model human placental development and susceptibility to emerging pathogens. Cell Stem Cell. 2022;29(5):810–25.e8.

22. Laukoetter MG, Nava P, Lee WY, Severson EA, Capaldo CT, Babbin BA, et al. JAM-A regulates permeability and inflammation in the intestine in vivo. J Exp Med. 2007;204(13):3067–76.

23. Li S, Edgar D, Fässler R, Wadsworth W, Yurchenco PD. The Role of Laminin in Embryonic Cell Polarization and Tissue Organization. Developmental Cell. 2003;4(5):613–24.

24. Schüler SC, Liu Y, Dumontier S, Grandbois M, Le Moal E, Cornelison D, et al. Extracellular matrix: Brick and mortar in the skeletal muscle stem cell niche. Front Cell Dev Biol. 2022;10:1056523.

25. Altavilla A, Manfredi C, Baiardi P, Dehlinger-Kremer M, Galletti P, Pozuelo AA, et al. Impact of the new european paediatric regulatory framework on ethics committees: overview and perspectives. Acta Paediatrica. 2012;101(1):e27–e32.

26. Parisi S, Piscitelli S, Passaro F, Russo T. HMGA Proteins in Stemness and Differentiation of Embryonic and Adult Stem Cells. Int J Mol Sci. 2020;21(1).

27. Hicks MR, Pyle AD. The emergence of the stem cell niche. Trends in Cell Biology. 2023;33(2):112–23.

28. Kordes C, Häussinger D. Hepatic stem cell niches. J Clin Invest. 2013;123(5):1874–80.

29. Pinho S, Frenette PS. Haematopoietic stem cell activity and interactions with the niche. Nature Reviews Molecular Cell Biology. 2019;20(5):303–20.

30. Beumer J, Clevers H. Cell fate specification and differentiation in the adult mammalian intestine. Nat Rev Mol Cell Biol. 2021;22(1):39–53.

31. Burton GJ, Jauniaux E. The human placenta: new perspectives on its formation and function during early pregnancy. Proc Biol Sci. 2023;290(1997):20230191.

32. Turco MY, Gardner L, Kay RG, Hamilton RS, Prater M, Hollinshead MS, et al. Trophoblast organoids as a model for maternal-fetal interactions during human placentation. Nature. 2018;564(7735):263-7.

33. Lee CQE, Turco MY, Gardner L, Simons BD, Hemberger M, Moffett A. Integrin α2 marks a niche of trophoblast progenitor cells in first trimester human placenta. Development. 2018;145(16).

34. Shannon MJ, McNeill GL, Koksal B, Baltayeva J, Wächter J, Castellana B, et al. Single-cell assessment of primary and stem cell-derived human trophoblast organoids as placenta- modeling platforms. Dev Cell. 2024;59(6):776–92.e11.

35. de Boer CJ, van Krieken JH, Janssen-van Rhijn CM, Litvinov SV. Expression of Ep-CAM in normal, regenerating, metaplastic, and neoplastic liver. J Pathol. 1999;188(2):201–6.

36. Schmelzer E, Wauthier E, Reid LM. The phenotypes of pluripotent human hepatic progenitors. Stem Cells. 2006;24(8):1852–8.

37. Zhang L, Theise N, Chua M, Reid LM. The stem cell niche of human livers: symmetry between development and regeneration. Hepatology. 2008;48(5):1598–607.

38. Sasamoto Y, Lee CAA, Wilson BJ, Buerger F, Martin G, Mishra A, et al. Limbal BCAM expression identifies a proliferative progenitor population capable of holoclone formation and corneal differentiation. Cell Rep. 2022;40(6):111166.

39. Wang X, Hallen NR, Lee M, Samuchiwal S, Ye Q, Buchheit KM, et al. Type 2 inflammation drives an airway basal stem cell program through insulin receptor substrate signaling. J Allergy Clin Immunol. 2023;151(6):1536–49.

40. Ragazzini R, Boeing S, Zanieri L, Green M, D’Agostino G, Bartolovic K, et al. Defining the identity and the niches of epithelial stem cells with highly pleiotropic multilineage potency in the human thymus. Developmental Cell. 2023;58(22):2428–46.e9.

41. Ellis SJ, Tanentzapf G. Integrin-mediated adhesion and stem-cell-niche interactions. Cell Tissue Res. 2010;339(1):121–30.

42. Chen S, Lewallen M, Xie T. Adhesion in the stem cell niche: biological roles and regulation. Development. 2013;140(2):255–65.

43. Fitzgerald B, Kingdom J, Keating S. Distal villous hypoplasia. Diagnostic Histopathology. 2012;18(5):195–200.

44. Benton SJ, Leavey K, Grynspan D, Cox BJ, Bainbridge SA. The clinical heterogeneity of preeclampsia is related to both placental gene expression and placental histopathology. Am J Obstet Gynecol. 2018;219(6):604.e1-.e25.

45. Farah O, Nguyen C, Tekkatte C, Parast MM. Trophoblast lineage-specific differentiation and associated alterations in preeclampsia and fetal growth restriction. Placenta. 2020;102:4–9.

46. Liu M, Liao L, Gao Y, Yin Y, Wei X, Xu Q, et al. BCAM Deficiency May Contribute to Preeclampsia by Suppressing the PIK3R6/p-STAT3 Signaling. Hypertension. 2022;79(12):2830–42.

47. Lane SW, Williams DA, Watt FM. Modulating the stem cell niche for tissue regeneration. Nat Biotechnol. 2014;32(8):795–803.

48. Rozo M, Li L, Fan CM. Targeting β1-integrin signaling enhances regeneration in aged and dystrophic muscle in mice. Nat Med. 2016;22(8):889–96.

49. Rayagiri SS, Ranaldi D, Raven A, Mohamad Azhar NIF, Lefebvre O, Zammit PS, et al. Basal lamina remodeling at the skeletal muscle stem cell niche mediates stem cell self-renewal. Nat Commun. 2018;9(1):1075.

50. Hao Y, Stuart T, Kowalski MH, Choudhary S, Hoffman P, Hartman A, et al. Dictionary learning for integrative, multimodal and scalable single-cell analysis. Nat Biotechnol. 2023.

51. Yu G, Wang LG, Han Y, He QY. clusterProfiler: an R package for comparing biological themes among gene clusters. Omics. 2012;16(5):284–7.

52. Jin S, Guerrero-Juarez CF, Zhang L, Chang I, Ramos R, Kuan C-H, et al. Inference and analysis of cell-cell communication using CellChat. Nature Communications. 2021;12(1):1088.

53. Garcia-Flores V, Xu Y, Pusod E, Romero R, Pique-Regi R, Gomez-Lopez N. Preparation of single-cell suspensions from the human placenta. Nat Protoc. 2023;18(3):732–54.

54. van Rijn B, Van Opstal D, van Koetsveld N, Knapen M, Gribnau J, Schäffers O. Generation of Trophoblast Organoids from Chorionic Villus Sampling. Organoids. 2024;3(1):54–66.

55. Dekkers JF, Alieva M, Wellens LM, Ariese HCR, Jamieson PR, Vonk AM, et al. High- resolution 3D imaging of fixed and cleared organoids. Nat Protoc. 2019;14(6):1756–71.

56. Doellinger J, Schneider A, Hoeller M, Lasch P. Sample Preparation by Easy Extraction and Digestion (SPEED) - A Universal, Rapid, and Detergent-free Protocol for Proteomics Based on Acid Extraction. Mol Cell Proteomics. 2020;19(1):209–22.

57. Johnston HE, Yadav K, Kirkpatrick JM, Biggs GS, Oxley D, Kramer HB, et al. Solvent Precipitation SP3 (SP4) Enhances Recovery for Proteomics Sample Preparation without Magnetic Beads. Analytical Chemistry. 2022;94(29):10320–8.

58. van Sluis M, Yu Q, van der Woude M, Gonzalo-Hansen C, Dealy SC, Janssens RC, et al. Transcription-coupled DNA–protein crosslink repair by CSB and CRL4CSA-mediated degradation. Nature Cell Biology. 2024;26(5):770–83.

